# Enzyme-Responsive DNA Condensates

**DOI:** 10.1101/2024.07.02.601714

**Authors:** Juliette Bucci, Layla Malouf, Diana A. Tanase, Nada Farag, Jacob R. Lamb, Serena Gentile, Erica Del Grosso, Clemens F. Kaminski, Lorenzo Di Michele, Francesco Ricci

## Abstract

Membrane-less compartments and organelles are widely acknowledged for their role in regulating cellular processes, and there is an urgent need to harness their full potential as both structural and functional elements of synthetic cells. Despite rapid progress, synthetically recapitulating the nonequilibrium, spatially distributed responses of natural membrane-less organelles remain elusive. Here we demonstrate that the activity of nucleic-acid cleaving enzymes can be localised within DNA-based membrane-less compartments by sequestering the respective DNA or RNA substrates. Reaction-diffusion processes lead to complex nonequilibrium patterns, dependent on enzyme concentration. By arresting similar dynamic patterns, we spatially organise different substrates in concentric sub-compartments, which can be then selectively addressed by different enzymes, demonstrating spatial distribution of enzymatic activity. Besides advancing our ability to engineer advanced biomimetic functions in synthetic membrane-less organelles, our results may facilitate the deployment of DNA-based condensates as microbioreactors or platforms for the detection and quantitation of enzymes and nucleic acids.

## INTRODUCTION

Living cells are the basic units of life, able to sustain the highly articulate and dynamic functionalities that underpin the emergent behaviours of all living systems. Despite their diversity and complexity, all cells share common features such as the ability to adapt, communicate, process information, grow and divide.^1-2^ Critical to these conserved functionalities are the compartmentalised architectures that separate a cell’s interior from the surrounding environment and maintain its internal heterogeneity. Most cellular pathways are indeed directly controlled by the flow of matter and information across membranes and cell walls, and by the compositional and physical diversity of organelles.^1,3^ For example, compartmentalisation is closely connected with enzymatic activity within the cell, which is delicately balanced through multi-level organisation and regulation.^4-5^ While cellular compartmentalisation often relies on proteolipid membranes, membrane-less compartments emerging from liquid-liquid phase separation or condensation of proteins and nucleic acids are increasingly recognised as ubiquitous and critical in regulating a variety of physiological and pathological processes.^6-8^ Indeed, *membrane-less organelles* found in eukaryotic cells, such as nucleoli, Cajal bodies and P-bodies, are involved in key pathways ranging from transcriptional and post-transcriptional regulation to ribosome biogenesis to RNA degradation.^9-12^

Recent years have seen tremendous efforts towards building synthetic cells, artificial devices that display basic features of living cells and show programmable and tuneable life-like functionalities, applicable in drug discovery, bioengineering and sensing.^13,14^ Like living cells, synthetic cells require compartments to contain their functional molecular machinery and regulate transport and communication with the outside environment. Membrane-based compartments, created from lipids^15-19^ or polymers^20,21^ constitute a common choice due to their similarity to cellular membranes. Membrane-less enclosures, constructed from synthetic biomolecular condensates, coacervates, or hydrogels, represent valuable alternatives to establish compartmentalisation,^22-26^ particularly for their ability to support dynamic behaviours inspired by biological membrane-less organelles.^27^

Owing to the programmability of base pairing, facile synthesis and functionalisation, and computational design tools^28-30^ nucleic acid nanotechnology has emerged as a valuable toolkit for engineering both structure and functionalities of synthetic cells.^31-33^ Specifically, synthetic DNA and RNA nanostructures, including branched DNA junctions^34-37^ and single-stranded DNA block copolymers,^38,39^ have been adopted to construct membrane-less compartments with advanced functionalities. These include the ability to capture and release molecular cargoes,^40,41^ host enzymatic reactions,^27,42-44^ interface with live cells,^45,46^ and undergo structural transformations induced by changes in pH,^47,48^ ionic conditions,^49^ or light.^40,50^ Advances have also been made towards establishing physical and chemical heterogeneity within DNA-based membrane-less compartments, for instance by exploiting reaction-diffusion processes^27^ or phase separation.^51-53.^ These non-homogeneous constructs imitate the internal sub-compartments observed in several classes of biological membrane-less organelles, including stress granules, nucleoli, L-granules and paraspeckles.^8,9,54-56^

Despite these advances, however, synthetic membrane-less compartments still largely lack the functional complexity of biological ones. Progress remains to be made when implementing dissipative biochemical processes in synthetic condensates, and program their spatiotemporal distribution within distinct, co-existing sub-compartments.

Here we demonstrate that DNA-based membrane-less synthetic cells can be engineered to localise the activity of different DNA repair enzymes, targeting specific nucleic-acid substrates. Exploiting reaction-diffusion mechanisms controlled by the relative size, binding affinity and concentration of the nucleic acid substrates and enzymes, we show that the synthetic cells can be engineered to sustain complex, nonequilibrium patterns that evolve in space and time (Figure 1a). Finally, we use reaction-diffusion processes^27^ to establish static sub-compartments in condensates which can be selectively and individually targeted by enzymes, hence demonstrating spatial control over enzymatic activity reminiscent of that observed in natural membrane-less organelles (Figure 1b). ^57^

**Figure 1.**
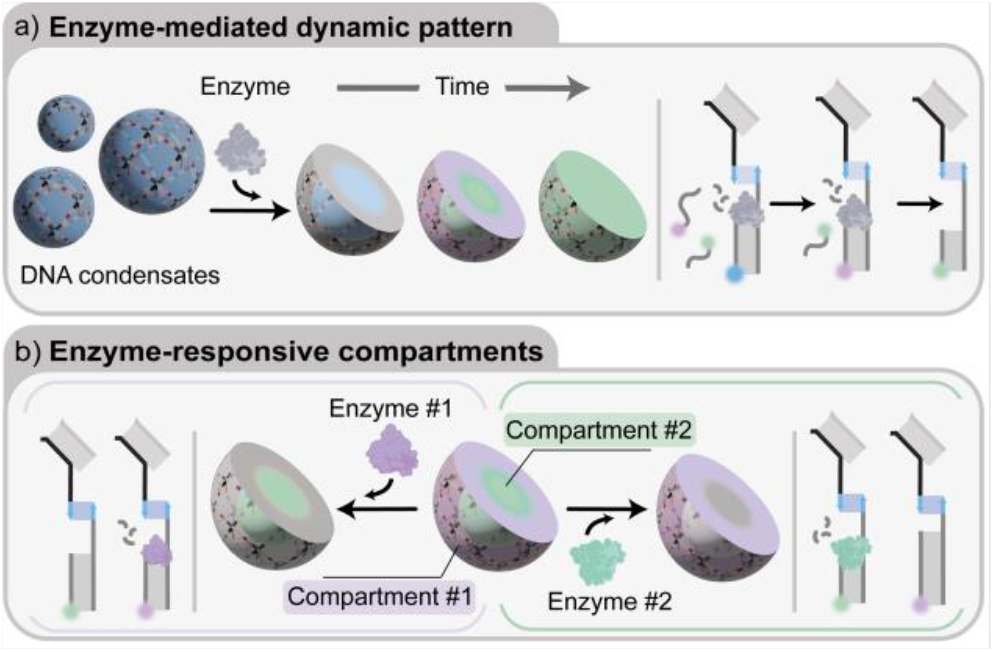
Enzyme-responsive DNA condensates. a) Schematic representation of enzyme-mediated dynamic patterning of DNA condensates. When added, an enzyme digests nucleic acid substrates bound to the condensates. Digested substrates can be replaced by fresh strands present in solution, producing reaction-diffusion patterns dependent on enzymatic activity, the diffusivity and binding strength of multiple, co-existing substrates. b) Schematic representation of DNA condensates with two concentric compartments each containing a different enzymatic substrate, allowing enzymatic reactions to occur only within the pre-determined sub-compartments.

## RESULT AND DISCUSSION

We constructed synthetic DNA condensates from the self-assembly of tetravalent DNA “nanostars”. As sketched in Figure 2a, individual nanostars fold from four distinct core strands forming a locked four-way DNA junction with 35 base-pair (bp) double-stranded DNA (dsDNA) arms. Each core strand is connected, through hybridisation of a 14-nucleotide (nt) domain, to a sticky strand terminating in a 10-nt “sticky end”. Nanostar-nanostar interaction are mediates by hybridisation of complementary sticky ends *α* and *α’*. Each nanostar also displays a 14-nt ssDNA domain (light blue), to which a 54-nt “anchor” strand (blue and orange) can be linked. The anchor strand serves as an addressable binding site for the enzyme-responsive moieties discussed below.

**Figure 2.**
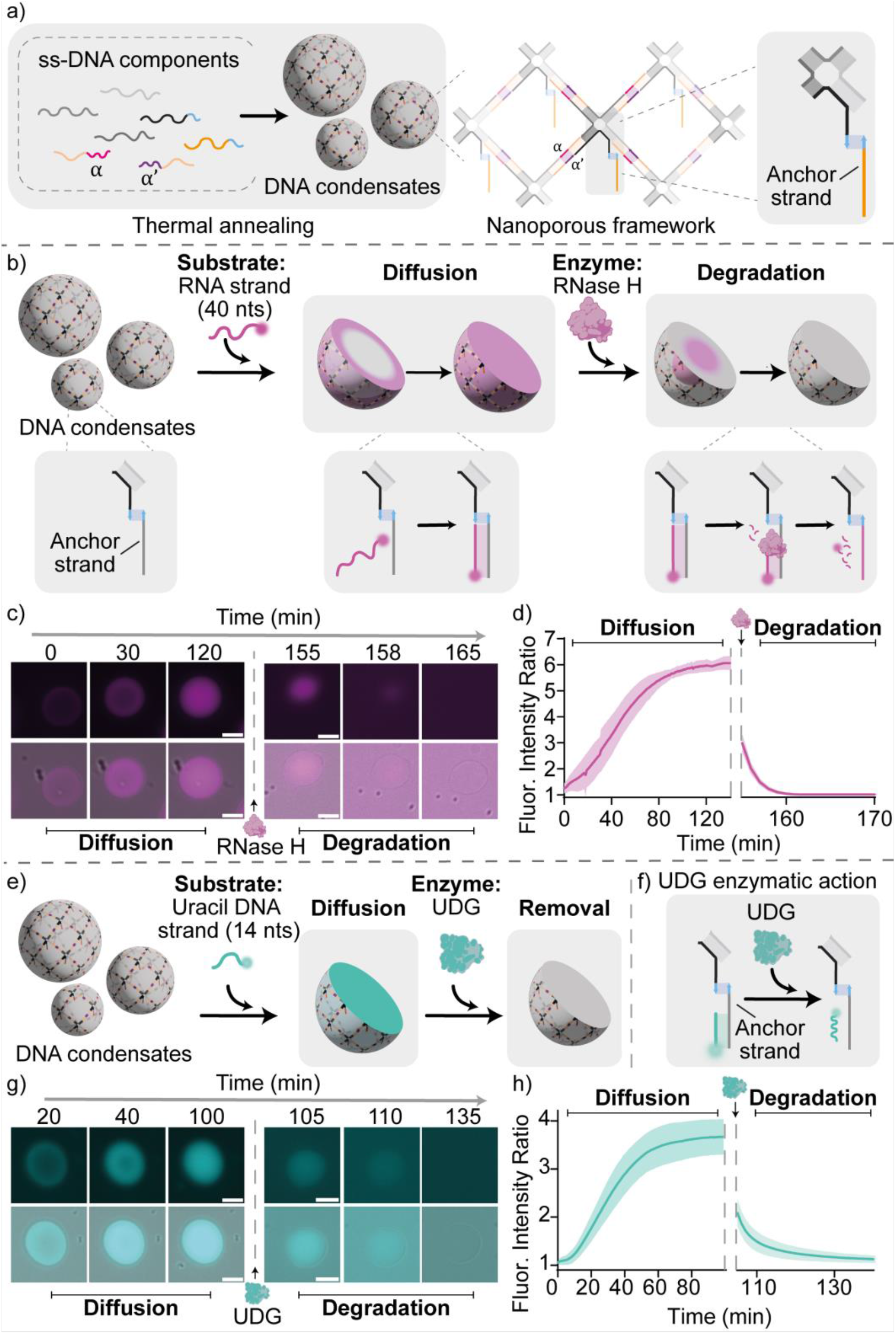
Enzyme-responsive DNA condensates. a) Nanoporous DNA condensates, hosting homogeneously distributed anchor strands, are obtained through slow thermal annealing from 90°C to 20°C of the ssDNA components. b) Cartoons and reaction schemes illustrating the diffusion and binding of a fluorophore-labelled RNA substrate within a DNA condensate, and its subsequent enzymatic degradation by RNase H. c) Epifluorescence micrographs (top) overlayed with brightfield images (bottom) of the diffusion, binding, and degradation process over time. d) Diffusion/binding and degradation kinetics tracked via the ratio of fluorescent signal samples within the condensates and the surrounding background. Data are shown as mean (solid line) ± standard deviation as obtained analysing n = 352/219 condensates (respectively diffusion stage/degradation stage) imaged across 3 technical replicates. f) Cartoons and reactions schemes illustrating the diffusion and binding of a fluorophore-labelled uracil DNA substrate and its degradation by UDG. g) Epifluorescence micrographs (top) overlayed with brightfield images (bottom) of the diffusion, binding, and degradation process over time. h) Diffusion/binding and degradation kinetics tracked via fluorescence intensity as for panel d. Data are shown as mean (solid line) ± standard deviation as obtained analysing n = 807/155 condensates (respectively diffusion stage/degradation stage) imaged across 12/3 technical replicates (respectively diffusion stage/degradation stage). Experiments were performed in Tris HCl 20 mM, EDTA 1 mM, MgCl_2_ 10 mM and 0.05 M NaCl; pH 8.0 at T=30°C. Sample preparation, annealing process and image analysis details are provided in SI Methods. All scale bars are 10 µm.

Nanostar folding and self-assembly are induced through the one-pot annealing of a stoichiometric mixture of the constituent strands (core strands, anchor strand, sticky strands), from 90°C to 20°C, leading to the formation of spherical condensate droplets with diameters ranging from 10 μm to 40 μm (Figure 2a, Figure S1). The distance between the junctions of two connected nanostars is ∼26 nm (80 bp), ensures a high porosity of the resulting network given the rigidity of dsDNA (persistence length ∼50nm^58^). In particular, we expect the pore size to enable internal diffusion of macromolecular solutes such as oligonucleotides and proteins.^53,59-61^

Having established nanoporous, addressable DNA condensates, we proceed to demonstrating the localisation of enzymatic activity within them. We initially focus on the endonuclease RNase H, an enzyme that degrades RNA only when hybridised to form an RNA/DNA heteroduplex.^62^ As the target enzymatic substrate for RNase H, we introduce an RNA strand (40 nucleotides (nt), light magenta) fully complementary to the anchor strand. The RNA strand is labelled with a fluorophore (Atto 488) so that its localisation within the condensates, and subsequent enzymatic degradation, can be monitored through epifluorescence imaging. Upon the addition of the RNA strand (200 nM) to a solution containing previously formed DNA condensates (200 nM of the DNA nanostars and anchor strands), we observe an inward-propagating front resulting from the diffusion of the RNA substrates through the DNA condensate, and their subsequent binding to the available DNA anchor strand. The diffusion transient is tracked in Figure 2c-d (left) by monitoring the mean fluorescence intensity within the condensate, determined through image segmentation as outlined in the SI Methods (Section *Image analysis*). A homogenous distribution of the RNA substrate through the condensate is achieved within 100 minutes, as demonstrated in Figure 2c,d (left) and in Movie S1. We then added RNase H (50 U/mL, magenta) to the RNA-loaded DNA condensates, observing the propagation of a degradation wave through the condensate as a result of the enzymatic reaction catalysed by RNase H (Figure 2c-d (right) and Movie S2). The degradation proceeds very rapidly, and the substrate strand is fully degraded within 10 minutes, as demonstrated in Figure 2d (right). A quick degradation is consistent with the observation that the enzyme has a hydrodynamic diameter of 2.2 nm,^63^ smaller than the estimated mesh size of the DNA networks and enabling rapid diffusion across the condensates. We stress that the enzymatic activity of RNase H does not affect the structure of the DNA condensates, which remain unchanged over the duration of the experiment (Figure S2).

As summarised in Figure S3 and S4, we demonstrated condensate-loading and subsequent digestion for RNA substrates of different length (Movie S3, S4). The three different RNA strands propagate at different rates through the condensate owing to the difference in size and hence diffusion constant,^64^ but are degraded by RNase H, when bound to DNA, at similar rates (Movie S5, S6).

The same approach for localising catalytic activity within the condensates can be extended to different enzymes, with the simple expedient of replacing the substrate bound to the anchor strand. To demonstrate this degree of versatility, we used uracil-DNA glycosylase (UDG, hydrodynamic diameter 5.8 nm),^63^ a base-excision repair enzyme that hydrolyses deoxyuridine mutations in DNA strands, leading to the formation of abasic sites.^65^ As the enzymatic substrate, we employed a fluorophore-labelled (Atto 488) 14 nt DNA strand complementary to the anchor strand and containing 4 deoxyuridine mutations (Figure 2e, f). Condensate loading was performed as discussed above, and shown in Figure 2g, h (left) and Movie S7. Upon the addition of UDG (25 U/mL), the enzymatic reaction induces the formation of apurinic sites that destabilise the duplex between the anchor strand and the substrate strand leading to the spontaneous dissociation of the latter. As a result, we observe a dissociation wave that completes within about 20 min (Figure 2 g, h right and Movie S8) which also does not structurally affect the DNA condensates (Figure S5).

Loading and subsequent digestion of the nucleic-acid substrates occur over different timescales, controlled by the diffusion rates of the macromolecules through the nanoporous condensates, the rates of hybridisation to the anchors strand, and the rates of enzymatic digestion. Some of these timescales are controllable by design, *e*.*g*. by changing the length of the nucleic-acid substrates or enzyme concentration. The interplay between these timescales can be exploited to establish dynamic compartments, whose composition evolves over time imitating the dynamic character of natural membrane-less organelles.

To demonstrate this functionality, we first exposed the DNA condensates to a mixture of three RNA substrate strands of different length and thus affinity for the anchor strand: 14 nt (Atto 550, blue), 25 nt (Atto 647, yellow), and 40 nt (Atto 488, magenta) (Figure S6). Upon addition of these three strands (200 nM each) to a solution containing the DNA condensates (200 nM of DNA nanostars and of anchor strands) we observe a time-dependent reaction-diffusion pattern, previously reported for DNA strands.^27^ The short (14 nt) strand enters the condensate first owing to faster diffusion, occupying the free anchor strands. The intermediate-length and diffusivity strand (25 nt) follows, displacing the short strand through a toehold-mediated strand displacement reaction.^66,67^ Finally, the longest and slowest-diffusing strand (40 nt) displaces the 25 nt strand, occupying all available anchor strands (Figure S6 b, c and Movie S9). This endpoint, depicted in Figure 3a (left), corresponds to thermodynamic equilibrium, given the much more negative hybridisation free energy of the 40 nt strand compared to the others.

**Figure 3.**
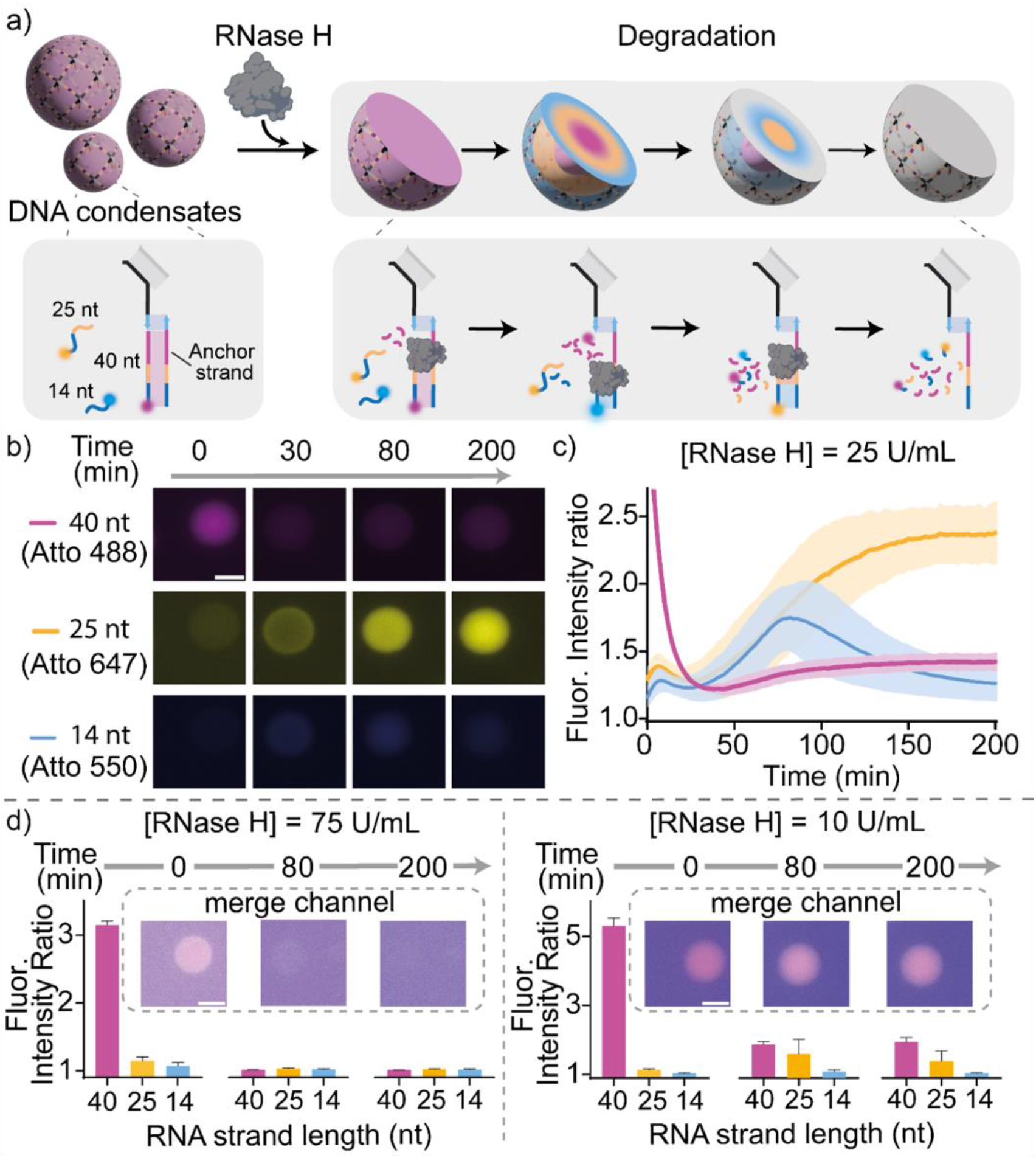
RNase H dynamic compartmentalisation. a) Cartoons and reaction schemes illustrating an expected degradation pattern induced by RNase H in the presence of three RNA substrate strands of different lengths (each labelled with a different fluorophore) competing for the anchor strand. b) Epifluorescence micrographs demonstrating the time-evolution of a typical condensate in a sample containing RNase H (25 U/mL) and the three RNA substrate strands (each at 200 nM). c) Ratio between the fluorescence intensity recorded within the condensate and the surrounding background for the three fluorescent constructs in samples corresponding to the experiment in panel b. Data are shown as mean (solid line) ± standard deviation as obtained analysing n = 195 condensates imaged across 3 technical replicates. d) Fluorescence intensity ratios (as in panel c) and epifluorescence micrographs at different times obtained using a fixed concentration of RNA strands (200 nM) and two different RNase H concentrations: 75 U/ml (left) and 10 U/mL (right). Data are shown as mean (solid line) ± standard deviation as obtained analysing n = 128/222 condensates (respectively RNase H 75 U/mL and 10 U/mL) imaged across 3 technical replicates. Experiments were performed in Tris HCl 20 mM, EDTA 1 mM, MgCl_2_ 10 mM and 0.05 M NaCl; pH 8.0 at T=30°C. Sample preparation, annealing process and image analysis details are provided in SI methods. All scale bars are 10 µm.

Upon adding RNase H, a dynamic reconfiguration is triggered within the condensates. The enzyme, which can only digest RNA when hybridised to DNA, first degrades the 40 nt RNA strands bound to the anchor strand. Removal of the strongest-binding strand allows the shorter strands to re-populate the condensates, as sketched in Figure 3a (right) and experimentally demonstrated in Figure 3b, c. Both the dynamics of the compartment-reconfiguration transient and its endpoint can be controlled by changing the concentrations of RNase H or of RNA strands. To demonstrate this, we carried out different reactions at a fixed concentration of DNA condensates (200 nM of the DNA nanostars and of anchor strands), in the presence of a mixture of substrate strands (each one at 200 nM) and at different concentrations of RNase H (75, 25, and 10 U/mL). At intermediate concentration of RNase H (25 U/mL, Figure 3b, c and Movie S10) we observe the rapid degradation of the longest RNA strand, followed by the two shorter strands rapidly occupying the binding sites made available, with the shortest being slightly faster. A transient configuration in which the two shortest RNA strands are homogenously distributed through the condensate thus emerge, which rapidly evolves due to the displacement of the shorter strand by the intermediate one (25 nts). Ultimately, the DNA condensates become mainly loaded with the intermediate strand, which remains indefinitely stable due to the progressive loss of activity of the enzyme (Figure 3b, d). Using a higher (75 U/mL) or lower (10 U/mL) concentration of RNase H results in distinctively different diffusion-degradation pathways, as demonstrated in Figure 3d by sampling the fluorescent signal from the three RNA substrates at three different times (0, 80 and 200 min). With [RNase H] = 75 U/mL, after the longest RNA is removed, the two others are degraded by the enzyme as soon as they bind the free anchor strands, and neither persists indefinitely. In turn, with 10 U/mL of RNase H, the longest RNA strand is only partially removed before loss of enzymatic activity, triggering occupation of some of the binding sites by the intermediate strand.

Having demonstrated the use of enzymatic reactions to program time-dependent responses, we proceed to show that activity can be localised in distinct, addressable sub-compartments, thus achieving spatial organisation of functionalities akin to that observed in biological condensates. As substrates for these reactions, we used two different nucleic acid strands: an RNA strand as the substrate of RNase H and an uracil-containing DNA strand as the substrate of UDG. Also, in this case the two substrate strands are labelled with two different fluorophores, so that their diffusion (and subsequent enzymatic removal) can be easily followed through fluorescence imaging. We proceeded to precisely localise these two nucleic acid substrates in distinct, concentric regions within condensates by employing a technique already optimised in a previous contribution and summarised in Figure S7.^27^ Specifically, adding the two strands in a solution containing the condensates results in a time-dependent reaction diffusion-pattern whereby the shorter (25 nt), faster-diffusing UDG substrate (Atto 488, cyan) initially diffuses and binds within the condensate, and is later displaced by the longer (40 nt), slower diffusing RNase H substrate (Atto 647, magenta) from the outside of the condensed inwards. A transient pattern is thus established, with the UDG substrate localised in the condensate’s core and the RNase H substrate in its outer shell. To arrest the pattern in this configuration, it is sufficient to add a large excess of a stopper strand (5 µM, 40 nt) with the same sequence of the binding site on the anchor strand, which captures all the free substrate strands present in solution. Once excess substrates are sequestered, the strand displacement reactions leading to pattern propagation are no longer possible, freezing the core-shell pattern in place and resulting in the formation of two concentric membrane-less compartments within the DNA condensates (Figure S7, 4a, Movie S11 and S12). We then proceeded to expose the patterned DNA condensates to one of, or both, the relevant enzymes, and to characterise the localised enzymatic activity with video microscopy and image analysis. We observe that enzymatic actions are mutually orthogonal and only occur within the target sub-compartments.

When only RNase H is added, enzymatic digestion is localised only in the external region (magenta) and the RNA strand is removed in about 15 min (Figure 4b top, Movie S13). If only UDG is added, the enzymatic activity is instead localised solely in the inner comportment, resulting in the removal of the uracil strand (Figure 4b middle, Movie S14). Volumetric reconstructions obtained through time-lapse Oblique Plane Microscopy (OPM) confirm the intended three-dimensional morphology of the patterned condensates, and the selective targeting of the outer shell and inner core when the condensates are exposed to RNase H or UDG, respectively (Figure 4b 3D view and Movie S15 and S16). OPM is a light sheet-based imaging modality that employs a single objective at the sample,^68^ allowing for the high spatial and temporal resolution afforded by light sheet imaging to be utilised with standard sample mounting technologies. Consequently, OPM is capable of imaging entire condensates at a temporal resolution unachievable by other 3D imaging technologies, such as laser scanning confocal microscopy. This provides an enhanced ability to study the three-dimensional dynamic response of the condensates upon the addition of the enzymes. If condensates are exposed to both UDG and RNase H, both substrate strands are removed (Figure 4b bottom, S8 and Movie S17).

**Figure 4.**
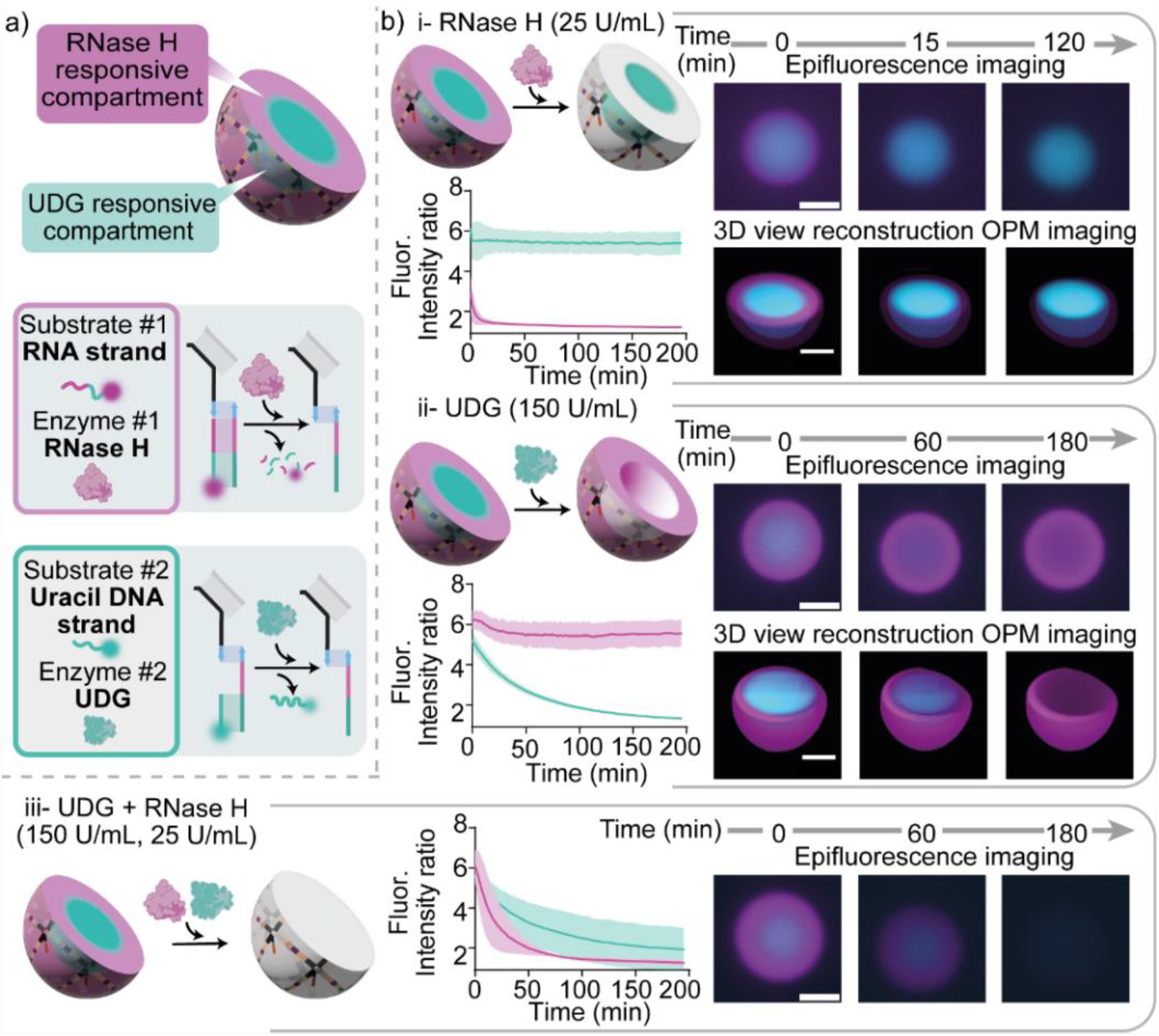
RNase H and UDG responsive compartments in DNA condensate. a) Cartoons illustrating the two responsive compartments in a DNA condensate: an external one (magenta) hosting the substrate of RNase H and an internal one (cyan) containing the substrate of UDG. b) Epifluorescence micrographs (top right), 3D reconstructions obtained from Oblique Plane Microscopy (bottom right) and Fluorescence intensity kinetics (left, as sampled with epifluorescence) demonstrating a localised, orthogonal and specific enzymatic activity within the condensates, by adding RNase H only (i), UDG only (ii) or both enzymes (iii). RNase H and UDG concentrations were fixed at 25 U/mL and 150 U/mL, respectively. RNA and uracil DNA substrates were fluorescently labelled with Atto 647 and Atto 488, respectively. Data are shown as mean (solid line) ± standard deviation as obtained analysing n = 115/93/91 condensates (respectively i, ii, iii) imaged across 3 technical replicates. Note that the sub-compartments established within the condensates do not change morphology over time, confirming that the condensates are in a solid phase and internal diffusion of the DNA nanostars (and anchor strands connected to them) does not occur over relevant experimental timescales. Experiments were performed in Tris HCl 20 mM, EDTA 1 mM, MgCl_2_ 10 mM and 0.05 M NaCl; pH 8.0 at T=30°C. Sample preparation, annealing process and image analysis details are provided in the SI Methods. All scale bars are 10 µm.

## CONCLUSION

Here we demonstrated that membrane-less, DNA-based condensates (synthetic cells), self-assembled from a small number of DNA and RNA oligonucleotides, can be engineered to sustain complex spatiotemporal patterns sustained by enzymatic reactions. The activity of DNA-repair enzymes RNase H and UDG is localised within the synthetic cells by hybridising their target nucleic-acid substrates to dedicated binding sites. The nano-porous nature of the condensates facilitates the diffusion of substrates, enzymes and products, enabling the emergence of complex non-equilibrium patterns that evolve in space and time, regulated by the relative size of the substrates, their affinity for the condensates and the concentration of the enzymes. Reaction-diffusion patterns can be arrested to precisely localise substrates in sub-compartments within the condensates, which can then be selectively targeted by the corresponding enzymes.

The nonequilibrium pattern formation and the spatial control of enzymatic activity are key characteristics of biological membrane-less organelles, and the ability to recapitulate them in artificial analogues could be highly valuable for the deployment of synthetic DNA condensates in bottom-up synthetic biology. For instance, one could consider exploiting enzyme-activity localisation to optimise enzymatic cascades hosted within synthetic cells, applicable to biomanufacturing and biosensing.^3,69-71^

In addition to expanding the temporal control over enzymatic activity with nucleic acids,^72-74^ the enzyme-dependent spatiotemporal patterns could be valuable as readouts for synthetic-cell based assays aimed at detecting substrates, enzymes and quantifying enzymatic activity, as relevant for diagnosing conditions characterised by dysregulation of enzymes or circulating nucleic acids. ^75-77^

## ASSOCIATED CONTENT

The Supporting Information is available free of charge via the Internet at http://pubs.acs.org.

Oligonucleotide sequences used, protocols, experimental methods, supplementary figures, image analysis, and supplementary videos description.

Movie S1: Diffusion substrate RNA strand 40 nt.

Movie S2: Degradation substrate RNA strand 40 nt in presence of 50 U/mL of RNase H.

Movie S3: Diffusion substrate RNA strand 14 nt. Movie S4: Diffusion substrate RNA strand 25 nt.

Movie S5: Degradation substrate RNA strand 14 nt in presence of 50 U/mL of RNase H.

Movie S6: Degradation substrate RNA strand 25 nt in presence of 50 U/mL of RNase H.

Movie S7: Diffusion substrate Uracil strand 14 nt.

Movie S8: Degradation substrate Uracil strand 14 nt of 25 U/mL of UDG.

Movie S9: Simultaneous diffusion of three RNA substrates with different length: 40 nt, 25 nt and 14 nt.

Movie S10: Dynamic pattern compartmentalisation with RNase H 25 U/mL

Movie S11: Formation of DNA condensates with core-shell compartments: Diffusion

Movie S12: Formation of DNA condensates with core-shell compartments: Stopper

Movie S13 DNA condensates with core-shell compartments exposed to RNase H.

Movie S14: DNA condensates with core-shell compartments exposed to UDG.

Movie S15: 3D view reconstruction of DNA condensates with core-shell compartments exposed to RNase H.

Movie S16: 3D view reconstruction of DNA condensates with core-shell compartments exposed to UDG.

Movie S17: DNA condensates with core-shell compartments exposed to both RNase H and UDG.

A dataset in support of this work will be made available on a public, permanent repository upon final publication. Prior to final publication, all data are available from the corresponding authors upon request.

## Notes

The authors declare no competing financial interest.

---

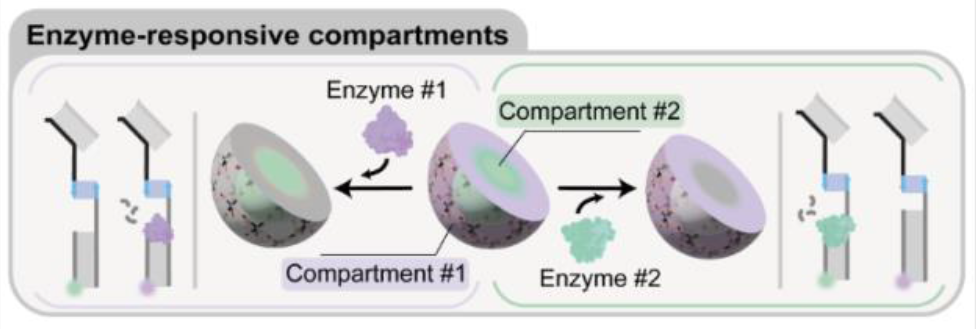

