## Supplementary Information for "Enzyme-Responsive DNA Condensates"

### Experimental Procedure

#### Materials

*Chemicals*

All reagent-grade chemicals, including DEPC-treated water, $MgCl_2$, Trizma hydrochloride, ethylenediaminetetraacetic acid (EDTA), NaCl, 1,4-Dithiothreitol (DTT), were purchased from Sigma-Aldrich and used without further purifications. 100× TE buffer was diluted in Milli-Q water prior to use, to an end concentration of 10 mM Tris, 1 mM EDTA, pH ~8.0. All buffer solutions were filtered through 0.22 µm syringe filters (Millex), stored at 4 °C and used within three weeks of preparation.

*Enzymes*

UDG and RNase H recombinant were purchased from New England Biolabs (Beverly, MA, USA).

*Oligonucleotides*

Oligonucleotides employed in this work were synthesised and purified using standard desalting for non-functionalised strand by Integrated DNA Technologies (IDT) and used without further purification. Fluorophore labelled oligonucleotides were HPLC-purified. DNA oligonucleotide strands were shipped lyophilised and reconstituted in 1xTE. The RNA oligonucleotides were dissolved in DEPC-treated water. All reconstituted oligonucleotide strands were stored at -20 °C until used. The name and the sequences of all the oligonucleotide strands used are listed below.

| **Name** | **Sequence** |
|---|---|
| Core1 | 5’-*CGA CGC CGT GAC GC*C GTG GCC TGT GAT TGA GGC GCT GCG TCG TCC ACC GTG TGA AAC TT**G TCC GTT CTA AAT C**-3’ |
| Core2 | 5’-*CGA CGC CGT GAC GC*G TTT CAC ACG GTG GAC GAC GCT CGG ACT AGA ACT GTC TCG AAC-3’ |
| Core3 | 5’-*CGA CGC CGT GAC GC*G TTC GAG ACA GTT CTA GTC CGT CGC GAA TAC GCC GTG CCG TGC-3’ |
| Core 4 | 5’-*CGA CGC CGT GAC GC*G CAC GGC ACG GCG TAT TCG CGT GCG CCT CAA TCA CAG GCC ACG-3’ |

| | |
|---|---|
| Sticky strand α | 5’- *GCG TCA CGG CGT CG*G CTG GCT GCG -3’ |
| Sticky strand α’ | 5’- *GCG TCA CGG CGT CG*C GCA GCC AGC -3’ |
| Anchor strand_1 | 5’-GGT GAG GTG AGT GGA GTG GAG GTG TAG GAG TGA GGG TAA GG**A TTT AGA ACG GAC**-3’ |
| RNA strand 40nt | 5’-CUU ACC CUC ACU CCU ACA CCU CCA CUC CAC UCA CCU CAC C-ATTO488-3’ |
| RNA strand 25nt | 5’-ACA CCU CCA CUC CAC UCA CCU CAC C-ATTO647-3' |
| RNA strand 14nt | 5'-CCA CUC ACC UCA CC-ATTO550-3' |
| Anchor strand_2 | 5’-GGA GCA GAT CAT GGA GTG GAG GTG TAG GAG TGA GGG TAA GG**A TTT AGA ACG GAC**-3’ |
| Uracil strand 14nt | 5'-CCA /ideoxyU/GA /ideoxyU/C/ideoxyU/ GC/ideoxyU/ CC-ATTO488-3' |
| Uracil strand 25nt pattern | 5’-ACA CC/ideoxyU/ CCA C/ideoxyU/C CA/ideoxyU/ GA/ideoxyU/ C/ideoxyU/G CTC C-ATTO488-3’ |
| RNA strand 40nt pattern | 5’-CUU ACC CUC ACU CCU ACA CCU CCA CUC CAC UCA CCU CAC C-ATTO647-3’ |
| Stopper strand | 5’-GGA GCA GAT CAT GGA GTG GAG GTG TAG GAG TGA GGG TAA G-3’ |

The Core strand 1-4 form the locked four-way DNA junction. The italicised bases represent the 14-nts hybridisation domain interacting with the sticky ends α and α’ (reported in italic) of the sticky strands. The bold bases in core 1 are complementary with the bold region of both the anchor strand. Anchor strand_1 was used to form DNA condensates able to bind RNA strand 40nt, 24nt and 14 nt. While Anchor strand_2 was used to form DNA condensates able to bind Uracil strand 14nt, Uracil strand 25nt pattern and RNA strand 40nt pattern.

#### DNA Condensate assembly

All condensates were annealed in rectangular glass capillaries of dimension 60mm × 4mm × 0.4mm (CMScientific) in a one pot self-assembly process. The glass capillary tubes were cleaned by ultrasonication in a solution of 1% Hellmanex III (Hellma) in deionised (DI) water for 30 min at 50 °C and then rinsed with DI water. The capillary tubes were then further ultrasonicated in 2-propanol (Sigma-Aldrich) at 40 °C for 30 minutes and dried under $N_2$ prior to use.

Single strand DNA components were mixed in the reaction buffer (Tris HCl 20 mM, EDTA 1 mM, $MgCl_2$ 10 mM and 0.05 M NaCl) in an Eppendorf tube. The core stands 1-4 (that formed the locked four-way DNA junction), the sticky strands α/α’ and the desired anchor strand were mixed at a ratio of 1:2:1. Thus the final concentration of the DNA junction is 2.5 µM, while the one for the sticky strands is 5 µM, for a total volume of 60 µL.
The mixture was pipetted into the cleaned glass capillary tubes. Then, both ends of the capillary were capped with equal amounts of mineral oil, and finally sealed to a glass coverslip using Araldite Rapid 2-component epoxy glue.
Samples were annealed using a Bio-Rad C1000 Touch thermal cycler with the following annealing protocol: Hold at 95 °C for 30 minutes, then cool from 85 °C to 50 °C at -0.04°C $min^{-1}$, then cool from 50 °C to room temperature at -0.5 °C $min^{-1}$.
Condensates were extracted from capillaries into an Eppendorf containing 60 µL of the reaction buffer for a minimum of 30 minutes. To extract the formed condensates, both ends of the capillary were cut open with a diamond tip pen.

#### Imaging Fluorescence experiments

Epifluorescence micrographs were obtained using a Nikon Eclipse Ti2-E inverted microscope, equipped with a digital camera (Hamamatsu ORCA-Flash4.0 V3), a tuneable light source (Lumencor SPECTRA X LED engine), and Plan Fluor 20× 0.75N.A and Plan Fluor 40× 0.95N.A dry objectives (Nikon). All samples were imaged in µ-Plate 384 Well Glass Optical Bottom (170 µm +/- 5 µm) and sealed with MicroAmp Optical Adhesive Film (ThermoFisher Scientific) to prevent evaporation during imaging.

*Enzyme-responsive DNA condensate experiments*

Reaction-diffusion experiment samples were prepared by pre-mixing the desired substrate (RNA strands or Uracil DNA strand) at a target concentration of 0.2 µM with the reaction buffer (Tris HCl 20 mM, EDTA 1 mM, $MgCl_2$ 10 mM and 0.05 M NaCl) in a µ-Plate 384 Well Glass Optical Bottom (ibidi). Previously prepared and extracted DNA condensates were then added to the mixture at a target concentration of base strands (or nanostars) equal to 0.2 µM, for a final reaction volume of 20 µL. Then, the well plate was sealed with an optical adhesive film and mounted on the microscopy stage with a built-in incubator previously heated at 30ºC. Imaging positions were previously selected to minimise the time elapsed from insertion of the condensates to the start of recording (carried out at 30ºC). When the diffusion process was completed, the desired enzyme was added in the wells at the desired concentration and then recording was started again. Experiments in which the stopper strand was added were conducted in the same way. After the desired diffusion pattern was reached (around 1h after the addition of the DNA condensates), the stopper strand was added at a final concentration of 5 μM, in high excess compared to the concentration of the patterning strands.

#### Image analysis

The Epifluorescence microscopy time series in Figure 2, 3 and 4, were analysed using custom code written in MATLAB R2019b+. Micrographs were exported as Tag Image File Format (.tiff) files automatically labelled by colour channel and timepoint. All image analysis was carried out on raw microscopy data, with no scaling or LUTs applied. The image processing steps for the various experiments are reported below:

The first step is to create an image mask containing all condensates in the image by first applying a gaussian filtering, and then by binarizing the filtered image. A background mask is then created by inverting the condensate mask. To ensure results were not biased by over-sampling, the background mask was then eroded using the MATLAB *imerode* function, using a disk-shaped structuring element. Both the condensate and background masks were then manually checked to ensure clusters were identified correctly.

The refined masks were applied to raw confocal images to determine the average fluorescence intensity for both the condensates and the background and used to calculate an average intensity ratio. This intensity ratio reflects the average fluorescence intensity in the condensates ($I_{cond}$) divided the average intensity in the background ($I_{back}$) phase. Data from this analysis are presented in Figure 2d,h, 3c,d and 4b and in Figure S2b, S3c,f, S4c,f, S5b and S6c.

#### Oblique Plane Microscopy Imaging

All Oblique Plane Microscopy (OPM) imaging in this work was performed on a custom-built system. A 1.2 NA 60X water immersion primary objective (UPLSAPO60XW) and a 0.95 NA air immersion secondary objective (UPLXAPO40X) were used with a custom Plossl configuration 203 mm focal length secondary tube lens, providing a remote refocus magnification of 1.33x. The remote space image was captured using a glass-tipped AMS-AGY v1 objective lens and imaged using a 200 mm focal length tube lens and three-channel colour splitter (Cairn Optosplit 3) onto a Photometrics Kinetix sCMOS camera. Imaging was performed at an angle of 35 degrees, giving an effective NA of 1.2 and 1.15 along the x and y axes of the sample, respectively.

All datasets were acquired using a three-channel colour splitter (Cairn Optosplit 3), allowing both domains to be imaged in parallel. Splitting the colour channels in this way extended the required camera ROI in the direction the channels were split. Therefore, to minimise the increased camera readout time due to requiring a larger

ROI, the camera was oriented so that the direction of pixel readout was parallel to the sheet width. Consequently, a total camera ROI of 1040 × 2400 pixels was used (1040 rows read out), giving an ROI per channel of 1040 × 800 pixels.

All datasets were acquired with an 80 μm wide sheet in the sample with a sheet NA of 0.15. In both experiments, a total lateral FOV of 100 μm was scanned using galvanometric mirror scanning in 201 images, giving a total imaged volume of 80 x 100 x 42 μm (xyz respectively). The total acquisition time per volume was approximately 20.1 seconds.

The sample was exposed to both 488 and 647 nm lasers simultaneously during acquisition, and a shutter (stepper motor controlled by an Arduino) was used to block the excitation beam path and prevent unnecessary light exposure between volumes. Within the colour splitter, fluorescence from the Atto 488 channel reflected off a 567 nm cut-on longpass dichroic mirror (Thorlabs DMLP567R) and was transmitted through a 525/39 emission filter (Thorlabs MF525-39). The Atto 647 channel was transmitted through both a 567 nm cut-on longpass dichroic mirror (Thorlabs DMLP567R) and a 650 nm cut-on longpass dichroic mirror (Thorlabs DMLP650R) before being transmitted through a 665 nm longpass emission filter (Thorlabs FGL665).

*RNase H and UDG responsive compartments in DNA condensate experiments*

Previously prepared and extracted DNA condensates were first core-shell patterned in a well plate, as described in section *Imaging Fluorescence experiments - Enzyme-responsive DNA condensate experiments.* Then the well plate was mounted on the OPM microscopy stage with an incubator previously heated at 30ºC. Imaging positions were previously selected to minimise the time elapsed from addition of the desired enzyme (carried out at 30ºC). The desired enzyme (UDG or RNase H) was added in the wells at the desired concentration and then recording was started.
Volumes were acquired every minute for 4 hours and every 2 minutes for 5 hours for the addition of RNase H and UDG experiments, respectively.

*OPM image analysis*

After the data was acquired, the raw frames were computationally separated into their respective colour channels. Once separated, the channels were manually aligned using features present in both channels (glass fragments). The datasets were then deskewed by a factor of 3.4, axially resampled by a factor of 2.4, and rotated to return the data to the original coordinate frame of the sample. Bilinear interpolation was employed for both operations.

To aid in the visualisation of the entire 3D structure of the condensate, the top half of the condensates is removed in the 3D projection views to allow for the structure of the inner domains to be visualised. The 3D projection views, obtained from OPM data, are presented in Figure 4b-i and ii.

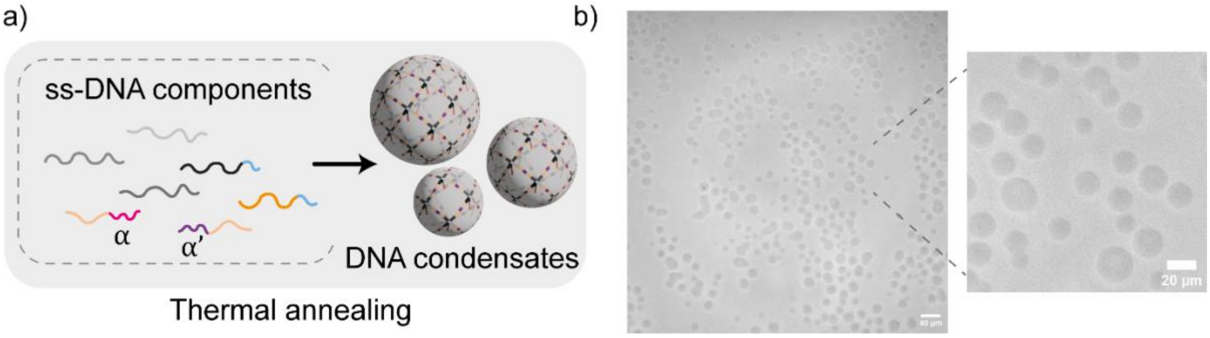


**Figure S1. DNA condensates formation.** a) Schematic illustrating the formation of the DNA condensates through slow thermal annealing from 90°C to 20 °C of the ssDNA components. b) Large field and zoomed in view of bright-field microscopy image showing the formed DNA condensates after the thermal annealing.

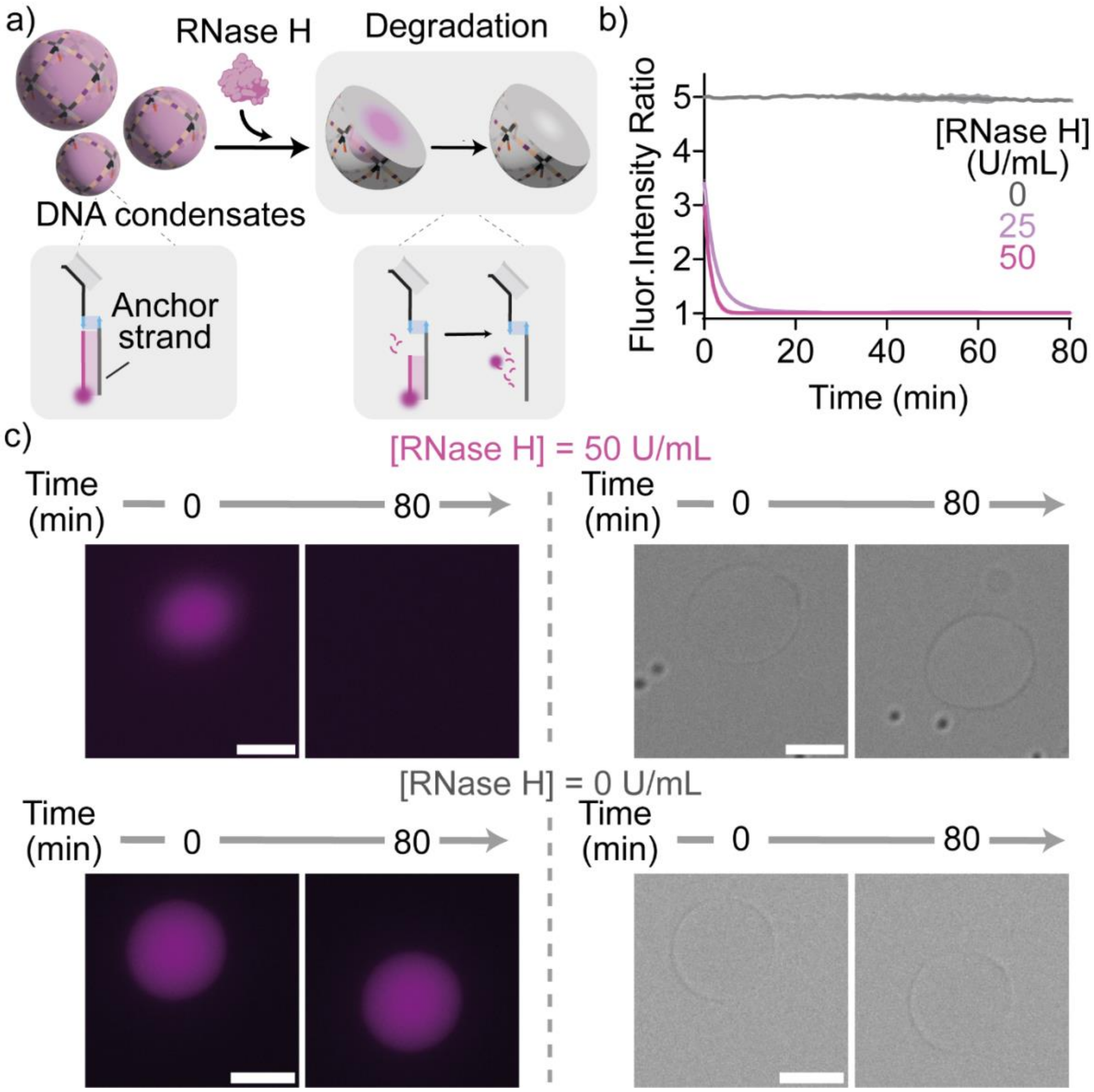


**Figure S2. RNase H-responsive DNA condensates using a 40 nt RNA substrate.** a) Cartoons and reactions schemes illustrating enzymatic degradation of a fluorophore-labelled (Atto 488, magenta) RNA substrate bound to the DNA condensates by RNase H. b) Degradation kinetics tracked *via* the ratio between the fluorescence intensity recorded within the condensates and the surrounding background, as extracted from epifluorescence images over time. In the absence of the enzyme (RNase H 0 U/mL, grey curve) the reaction does not proceed. Data are shown as mean (solid line) ± standard deviation as obtained analysing $n$ = 219/179/124 condensates (respectively degradation with 50, 25 and 0 U/mL of RNase H) imaged across 3/2 technical replicates (respectively 50, 25 U/mL of RNase H and 0 U/mL of RNase H). c) Epifluorescence (left) and bright-field (right) micrographs corresponding to the experiment in panel b. Experimental conditions used here are the same as in Figure 2. All scale bars are 10 µm.

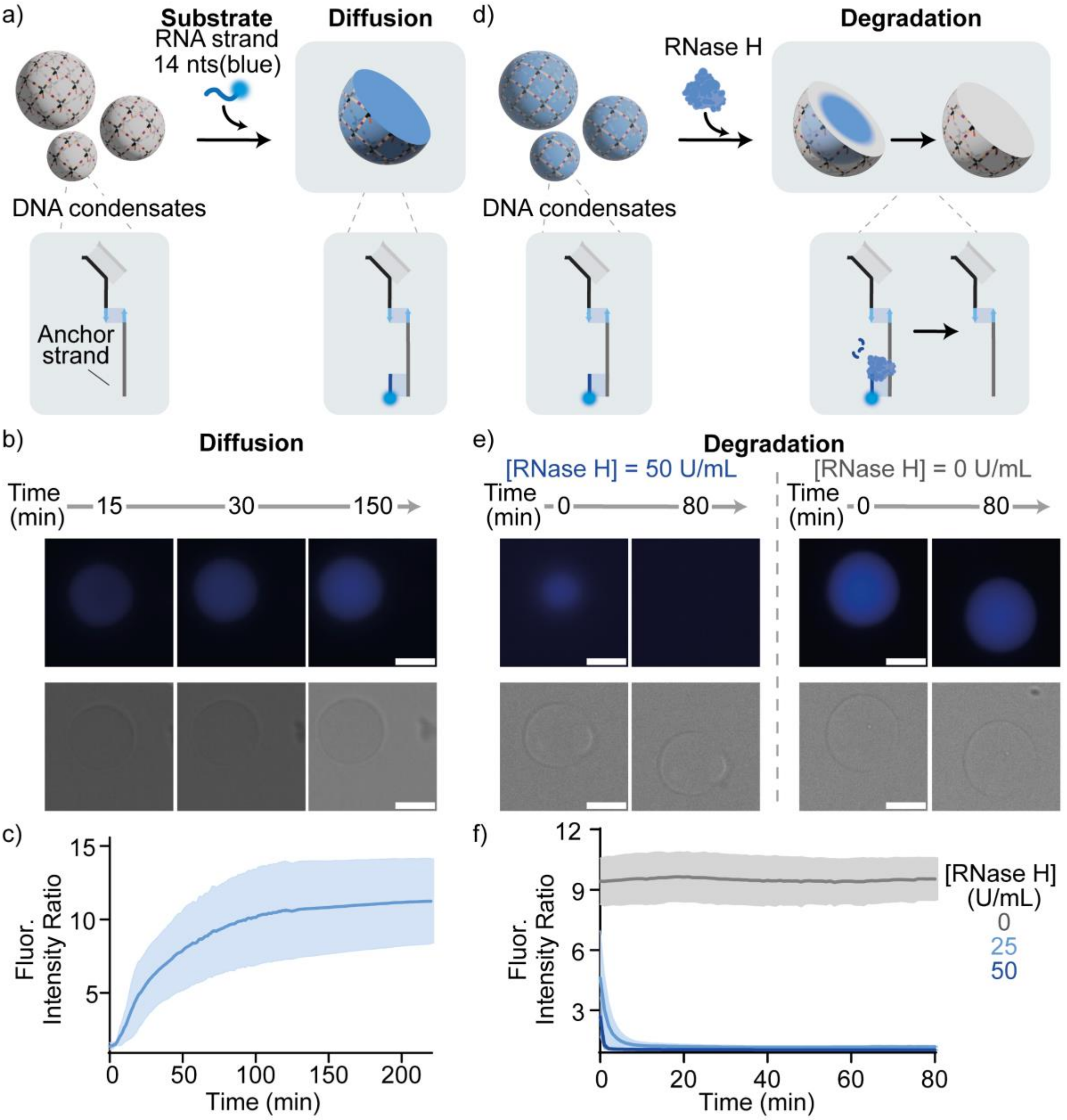


**Figure S3. RNase H-responsive DNA condensates using a 14 nt RNA substrate** a) Cartoons and reactions schemes illustrating diffusion and binding of a fluorophore-labelled (Atto 550, blue) 14 nt RNA substrate within DNA condensates. b) Epifluorescence (top) and bright field (bottom) micrographs of demonstrating the diffusion/binding process. c) Diffusion/binding kinetics tracked *via* the ratio between the fluorescence intensity recorded within the condensates and the surrounding background, as extracted from epifluorescence images over time. Data are shown as mean (solid line) ± standard deviation as obtained analysing $n$ = 295 condensates, imaged across 3 technical replicates. d) Cartoons and reactions schemes illustrating degradation of the substrate by RNase H. e) Epifluorescence (top) and bright field (bottom) micrographs of demonstrating the degradation process. f) Degradation kinetics monitored *via* fluorescence intensity as in panel c. Data are shown as mean (solid line) ± standard deviation as obtained analysing $n$ = 123/170/134 condensates (respectively degradation with 50, 25 and 0 U/mL of RNase H) imaged across 3/2 technical replicates (respectively for 50, 25 U/mL of RNase H and for 0 U/mL of RNase H). In the absence of the enzyme (RNase H 0 U/mL, grey curve) the reaction does not proceed. Experimental conditions used here are the same as in Figure 2. All scale bars are 10 µm.

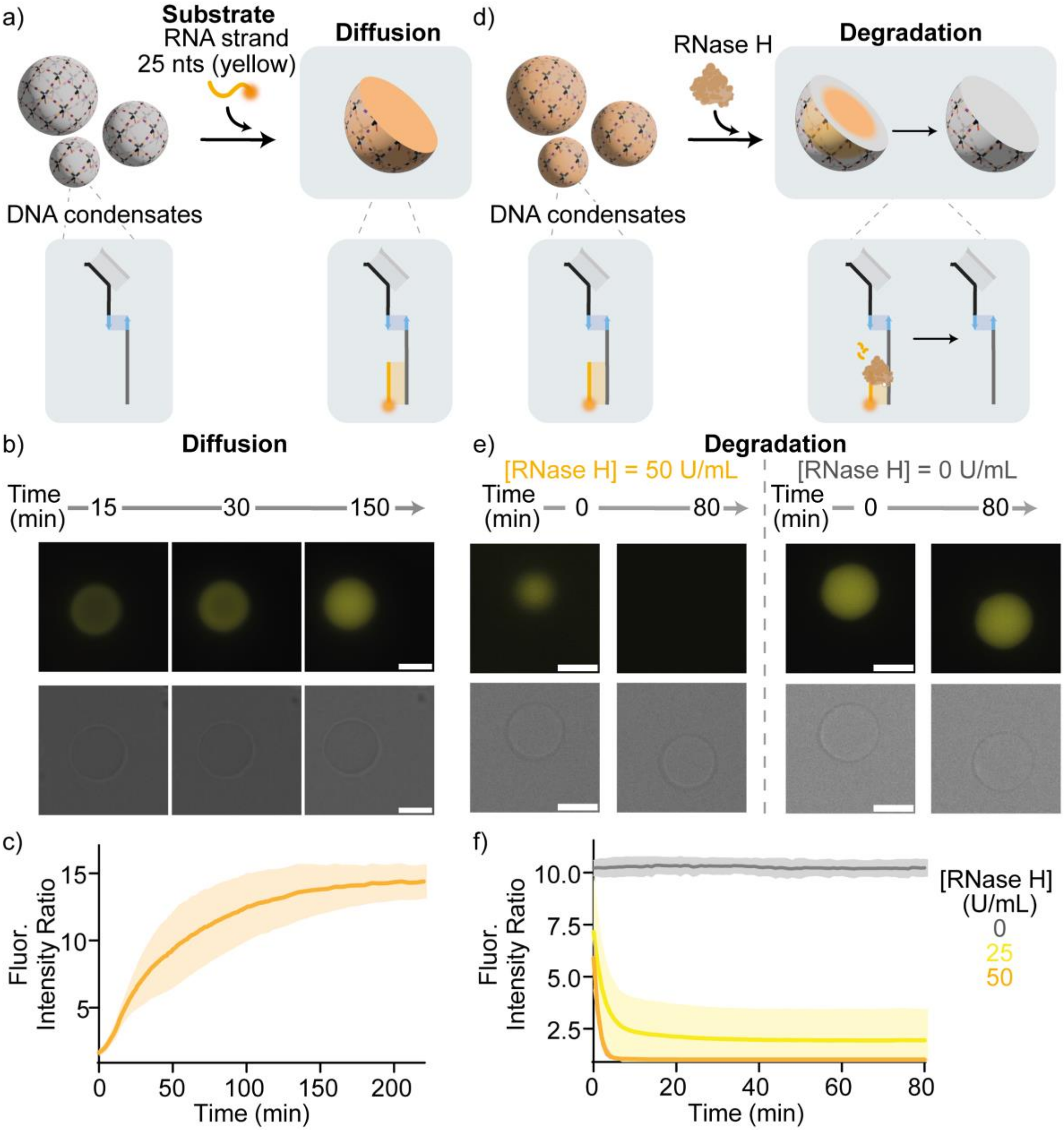


**Figure S4. RNase H -responsive DNA condensates using a 25 nt RNA substrate** a) Cartoons and reactions schemes illustrating diffusion and binding of a fluorophore-labelled (Atto 647, yellow) 25 nt RNA substrate within the condensates. b) Epifluorescence (top) and bright field (bottom) micrographs of demonstrating the diffusion/binding process. c) Diffusion/binding kinetics tracked *via* the ratio between the fluorescence intensity recorded within the condensates and the surrounding background, as extracted from epifluorescence images over time. Data are shown as mean (solid line) ± standard deviation as obtained analysing $n$ = 375 condensates, imaged across 4 technical replicates. d) Cartoons and reactions schemes illustrating degradation of the substrate by RNase H. e) Epifluorescence (top) and bright field (bottom) micrographs of demonstrating the degradation process. f) Degradation kinetics monitored *via* fluorescence intensity as in panel c. Data are shown as mean (solid line) ± standard deviation as obtained analysing $n$ = 149/163/144 condensates (respectively degradation with 50, 25 and 0 U/mL of RNase H) imaged across 3/2 technical replicates (respectively for 50, 25 U/mL of RNase H and for 0 U/mL of RNase H). In the absence of the enzyme (RNase H 0 U/mL, grey curve) the reaction does not proceed. Experimental conditions used here are the same as in Figure 2. All scale bars are 10 μm.

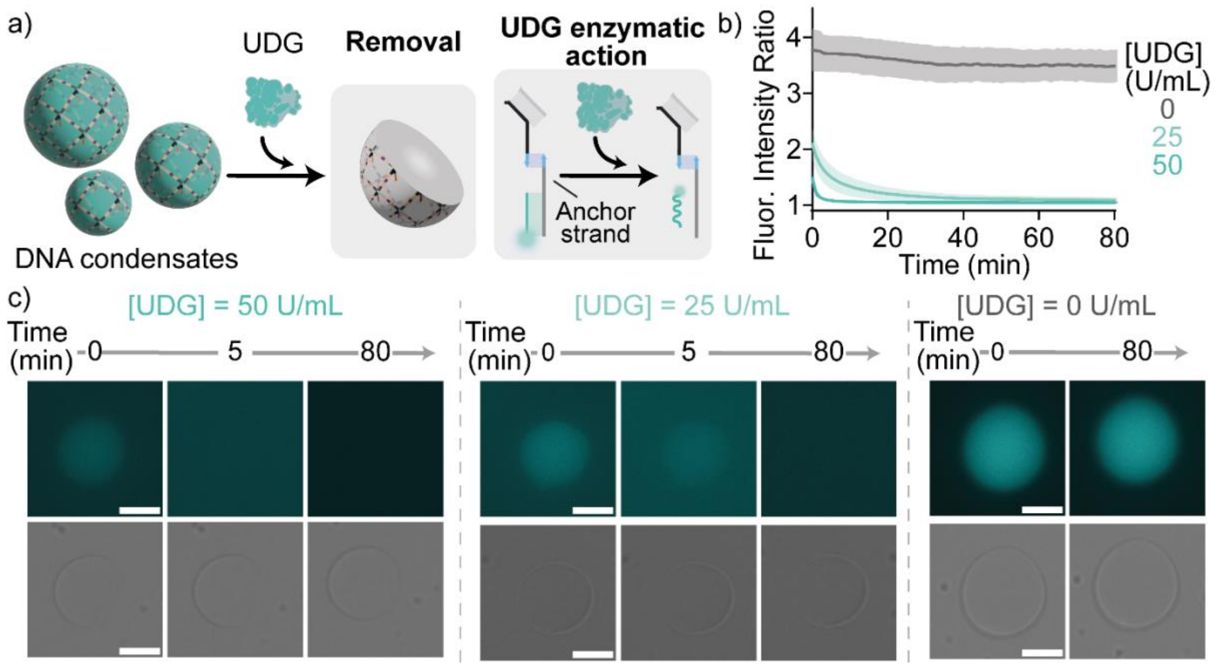


**Figure S5. UDG - responsive DNA condensates using a 14 nts DNA substrate**. a) Cartoons and reactions schemes illustrating degradation of uracil DNA substates by UDG within the condensates. b) degradation kinetics tracked *via* the ratio between the fluorescence intensity recorded within the condensates and the surrounding background, as extracted from epifluorescence images over time. Data are shown as mean (solid line) ± standard deviation as obtained analysing *n* = 105/155/160 condensates (respectively degradation with 50, 25 and 0 U/mL of UDG) imaged across 3 technical replicates. In the absence of the enzyme (UDG 0 U/mL, grey curve) the reaction does not proceed. c) Epifluorescence (top) and bright field (bottom) micrographs of demonstrating the degradation process. Experimental conditions used here are the same as in Figure 2. All scale bars are 10 µm.

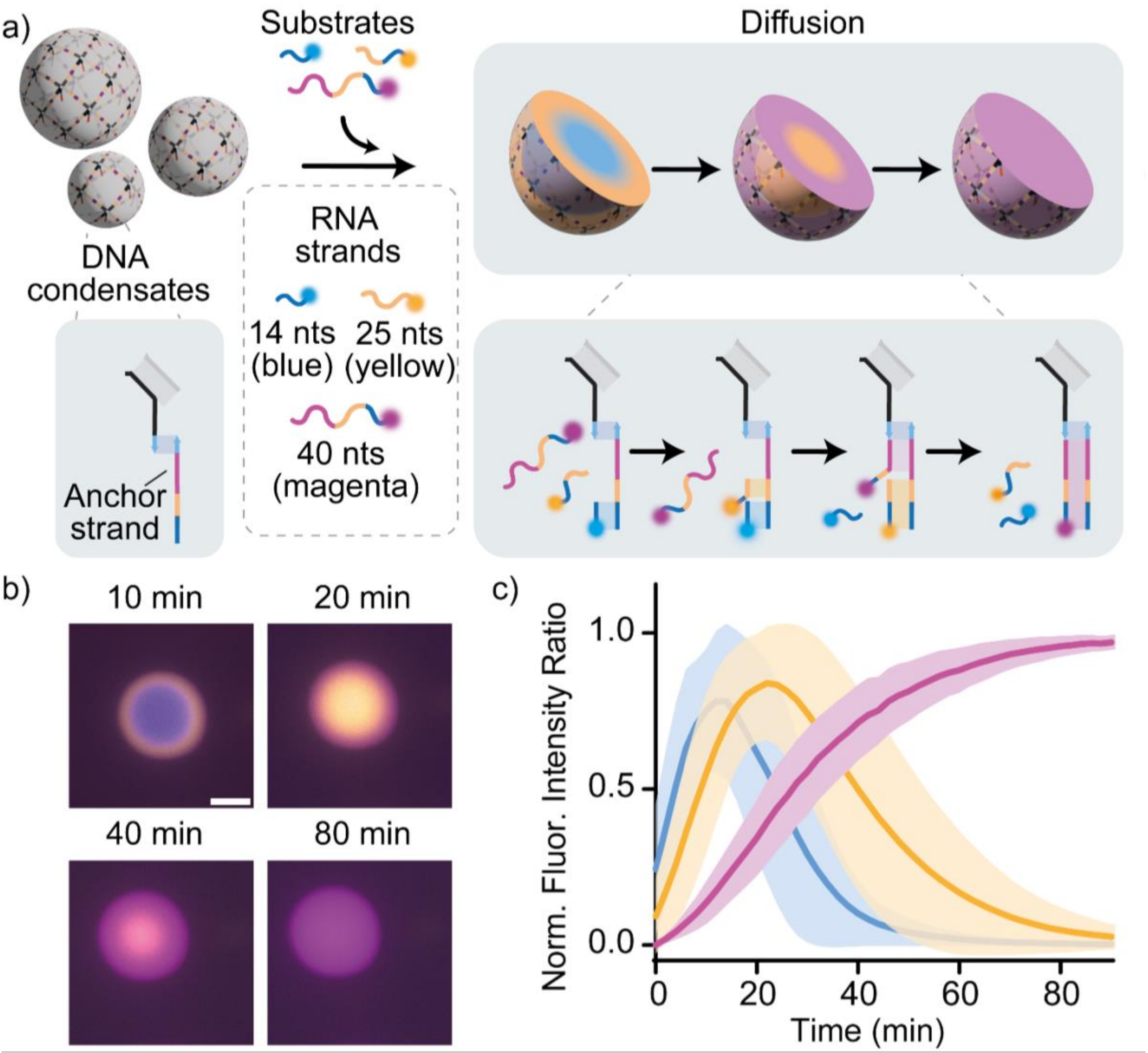


**Figure S6. RNA strand diffusion in DNA condensates.** a) Cartoons and reaction schemes illustrating the reaction-diffusion process originating when exposing non-functionalised DNA condensates (*i.e.* with free anchor strands) to three substrate RNA strands of different lengths, each labelled with a different fluorophore): 14 nt - Atto 550 (blue), 25 nt - Atto 647 (yellow), 40 nt - Atto 488 (Magenta). b) Epifluorescence images of the reaction diffusion process obtained at a fixed concentration of DNA condensates (200 nM of DNA nanostar and anchor strands) in presence of the three RNA strands (each at 200 nM). c) Reaction-diffusion transient tracked via the ratio between the fluorescence intensity recorded within the condensates and the surrounding background for each channel. To facilitate visualisation, curves are normalised with respect to their maximum intensity over the experimental time-window. As expected, the shortest strand diffuses first within the condensates, occupying free anchor sites. This is later displaced by the 25 nt strand, diffusing more slowly but binding more strongly. Finally, the longest, strongest binding strand occupies all binding sites. Data are shown as mean (solid line) ± standard deviation as obtained analysing $n$ = 873 condensates, imaged across 15 technical replicates. Experiments were performed in Tris HCl 20 mM, EDTA 1 mM, $MgCl_2$ 10 mM and 0.05 M NaCl; pH 8.0 at T=30°C. All scale bars are 10 µm.

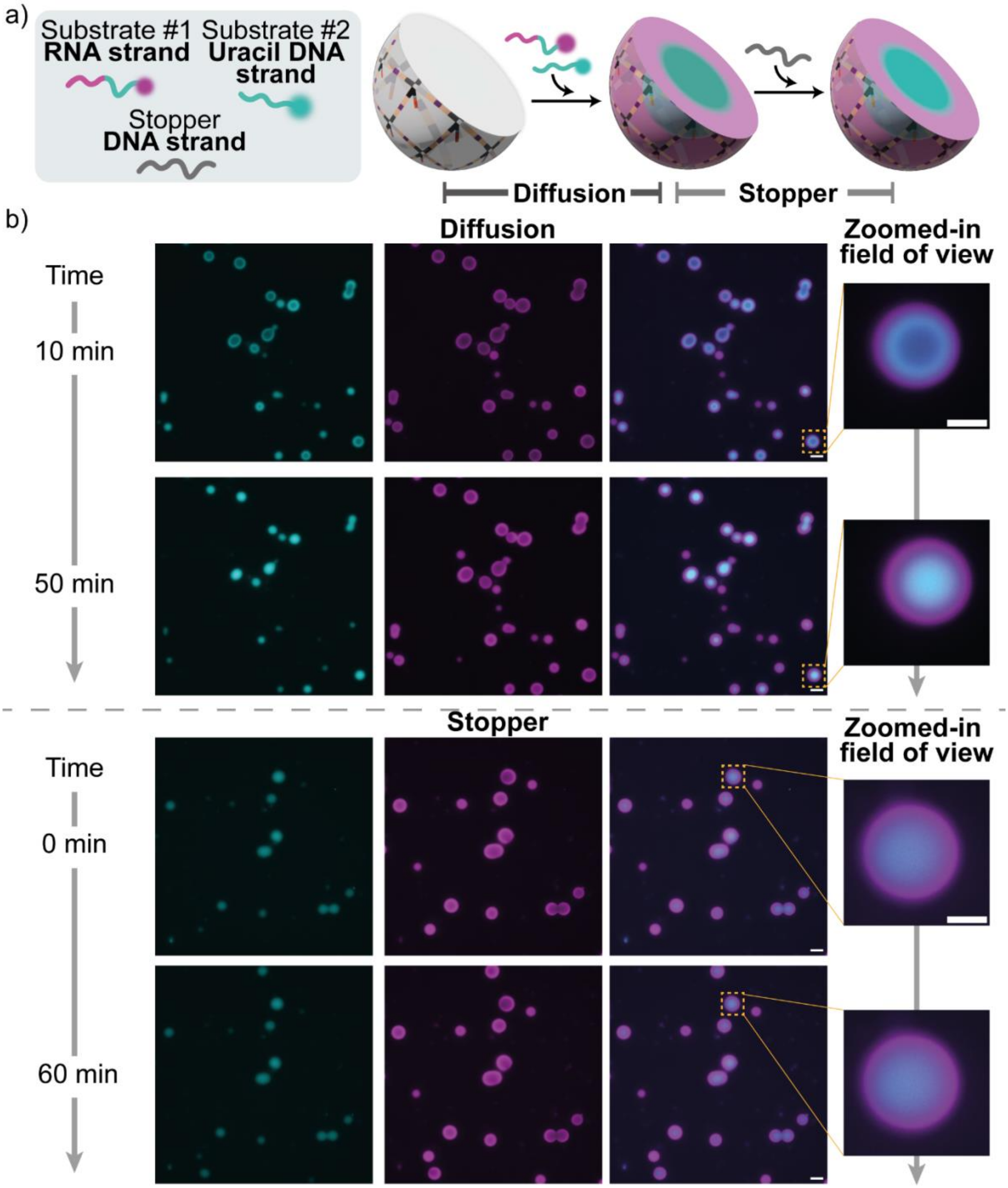


**Figure S7. Compartmentalisation of DNA condensates with substrate strands through core-shell patterning.** a) Cartoons and reaction schemes illustrating the reaction-diffusion process through which two substrate strands – the uracil DNA strand (25 nt, Atto 488 labelled, cyan) and the RNA strand (40 nt, Atto 647 labelled, magenta) – establish a core-shell pattern within the DNA condensates. Adding an excess of the stopper strand arrests pattern propagation by sequestering unbound substrate strands, resulting in the formation of two compartment. b) Epifluorescence images (left:large field of view and right: zoomed-in field of view) of the reaction-diffusion process and its subsequent stopping obtained at a fixed concentration of DNA condensates (200 nM DNA nanostar and of anchor strand), substrate strands (200 nM each) and adding the stop strand 5 µM). Note that arrested patters remain static over time, confirming that the condenses are in a solid phase. Experiments were performed in Tris HCl 20 mM, EDTA 1 mM, $MgCl_2$ 10 mM and 0.05 M NaCl; pH 8.0 at T=30°C. Scale bar 40 µm for large field of view and 10 µm for zoomed-in field of view.

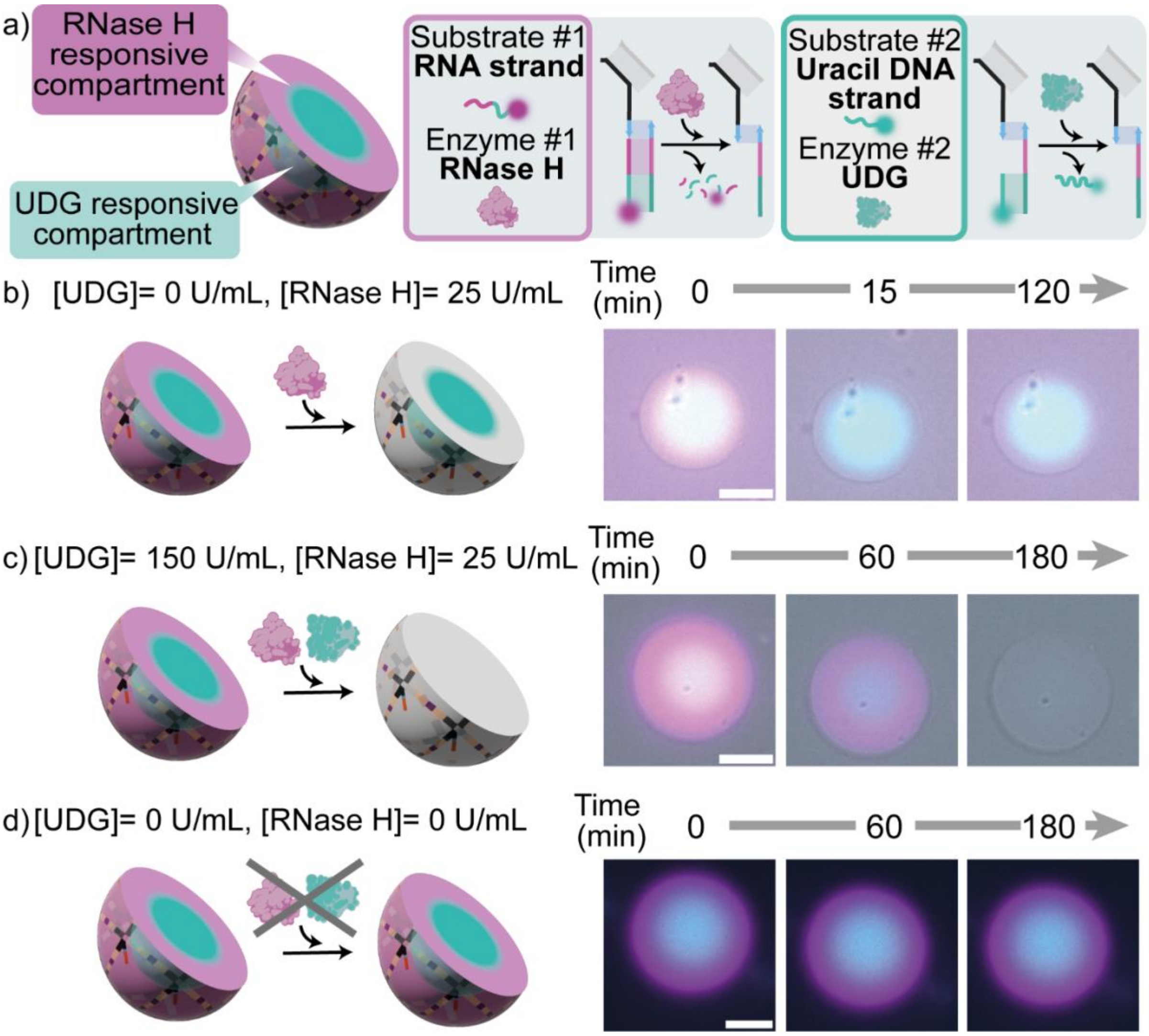


**Figure S8. RNase H and UDG responsive compartments in DNA condensate.** a) Cartoons illustrating the two responsive compartments in a DNA condensate: an external one (magenta) hosting the substrate of RNase H and an internal one (cyan) containing the substrate of UDG. b) Epifluorescence micrographs overlayed with brightfield images of condensates exposed only to RNase H. The substrate is removed from the shell without effecting the condensate structure overtime. c) Epifluorescence micrographs overlayed with brightfield images of condensates exposed to both UDG and RNase H. The core-shell pattern disappears, without effecting the condensate structure over time. d) Epifluorescence micrographs of condensates not exposed to enzymes. In the absence of the enzymes the patterned DNA condensate remain stable over time. Experimental conditions used here are the same as in Figure 4. All scale bars are 10 μm.

#### Supplementary movie description

**Movie S1:** Bright field and Epifluorescence time-lapse of a zoomed-in view of the diffusion and binding process of the substrate RNA strand (40 nts) within DNA condensates. The RNA strand is labeled with Atto 488 (shown in magenta). From the left: Bright field (BF), Atto 488, Merge: BF, Atto 488. Scale bar 10 µM.

**Movie S2:** Bright field and Epifluorescence time-lapse of a zoomed-in view of the substrate RNA strand 40 nt degradation within a DNA condensate, in presence of 50 U/mL of RNase H. The RNA strand is labeled with Atto 488 (shown in magenta). From the left: Bright field (BF), Atto 488, Merge: BF, Atto 488. Scale bar 10 µM.

**Movie S3:** Bright field and Epifluorescence time-lapse of a zoomed-in view of the diffusion and binding process of the substrate RNA strand (14 nt) within DNA condensates. The RNA strand is labeled with Atto 550 (shown in blue). From the left: Bright field (BF), Atto 550, Merge: BF, Atto 550. Scale bar 10 µM.

**Movie S4:** Bright field and Epifluorescence time-lapse of a zoomed-in view of the diffusion and binding process of the substrate RNA strand (25 nt) within DNA condensates. The RNA strand is labeled with Atto 647 (shown in yellow). From the left: Bright field (BF), Atto 647, Merge: BF, Atto 647. Scale bar 10 µM.

**Movie S5:** Bright field and Epifluorescence time-lapse of a zoomed-in view of the substrate RNA strand 14 nt degradation within a DNA condensate, in presence of 50 U/mL of RNase H. The RNA strand is labeled with Atto 488 (shown in blue). From the left: Bright field (BF), Atto 550, Merge: BF, Atto 550. Scale bar 10 µM.

**Movie S6:** Bright field and Epifluorescence time-lapse of a zoomed-in view of the substrate RNA strand 25 nt degradation within a DNA condensate, in presence of 50 U/mL of RNase H. The RNA strand is labeled with Atto 488 (shown in yellow). From the left: Bright field (BF), Atto 647, Merge: BF, Atto 647. Scale bar 10 µM.

**Movie S7:** Bright field and Epifluorescence time-lapse of a zoomed-in view of the diffusion and binding process of the substrate Uracil strand 14 nts degradation within DNA condensates. The Uracil strand is labeled with Atto 488 (shown in cyan). From the left: Bright field (BF), Atto 488, Merge: BF, Atto 488. Scale bar 10 µM.

**Movie S8:** Bright field and Epifluorescence time-lapse of a zoomed-in view of the substrate Uracil strand 14 nt degradation within a DNA condensate, in presence of 25 U/mL of UDG. The Uracil strand is labeled with Atto 488 (shown in cyan). From the left: Bright field (BF), Atto 488, Merge: BF, Atto 488. Scale bar 10 µM.

**Movie S9:** Bright field and Epifluorescence time-lapse of a zoomed-in view of the diffusion and binding process of three RNA strand (40, 25 and 14 nts, respectively magenta, yellow and blue) within DNA condensates. From the left: BF, Atto 488, Atto 647, Atto 550, Merge: Atto 488, Atto 647 Atto 550, Merge: BF, Atto 488 , Atto 647, Atto 550. Scale bar 10 μM.

**Movie S10:** Bright field and Epifluorescence time-lapse of a zoomed-in view of the dynamic patterning in presence of 10 U/mL of RNase H. From the left: BF, Atto 488, Atto 647, Atto 550, Merge: Atto 488, Atto 647 Atto 550, Merge: BF, Atto 488, Atto 647, Atto 550. Scale bar 10 μM.

**Movie S11:** Bright field and Epifluorescence time-lapse of a zoomed-in view of the formation of core-shell pattern DNA condensates: Diffusion and binding process of substrate RNA strand (40 nt) and substrate Uracil strand (25 nt), respectively magenta and cyan, within DNA condensates. From the left: bright field, Atto 488, Atto 647, Merge Atto 488 and 647, Merge BF, Atto 488 and Atto 647. Scale bar 10 μM.

**Movie S12:** Bright field and Epifluorescence time-lapse of a zoomed-in view of the formation of core-shell pattern DNA condensates: Stopper. The diffusion and binding of substrate RNA strand (magenta) and substrate Uracil strand (cyan) is stopped by the addition of the stopper strand. From the left: bright field, Atto 488, Atto 647, Merge: Atto 488,647, Merge: BF Atto 488, 647. Scale bar 10 μM.

**Movie S13:** Bright field and Epifluorescence time-lapse of a zoomed-in view of core-shell pattern DNA condensate exposed to RNase H (25 U/mL). From the left: BF, Atto 488, Atto 647, Merge: Atto 488,647, Merge: BF, Atto 488, Atto 647. Scale bar 10 μM.

**Movie S14:** Bright field and Epifluorescence time-lapse of a zoomed-in view of a core-shell pattern DNA condensates exposed to UDG (150 U/mL). From the left: BF, Atto 488, Atto 647, Merge: Atto 488,647, Merge: BF, Atto 488, Atto 647. Scale bar 10 μM.

**Movie S15:** A video illustrating the 3D view reconstruction a core-shell DNA condensates exposed to RNase H (25 U/mL). From the left: Atto 647, Atto 488, Merge: Atto 647, Atto 488. Scale bar 10 μM.

**Movie S16:** A video illustrating the 3D view reconstruction a core-shell DNA condensates exposed to UDG (150 U/mL). From the left: Atto 647, Atto 488, Merge: Atto 647, Atto 488. Scale bar 10 μM.

**Movie S17:** Bright field and Epifluorescence time-lapse of a zoomed-in view of a core-shell pattern DNA condensates exposed to both UDG (150 U/mL) and RNase H (25 U/mL). From the left: BF, Atto 488, Atto 647, Merge: Atto 488, Atto 647, Merge: BF, Atto 488, Atto 647. Scale bar 10 μM.
